# Transepithelial/endothelial electrical measurement using a low-cost and customizable Arduino-based sensor

**DOI:** 10.1101/2025.02.16.638578

**Authors:** Davide Lattanzi, Ludovica Montesi, Alice Sartini, Rossana Rauti

## Abstract

Transepithelial/transendothelial electrical resistance (TEER) is a label-free assay that is commonly used to assess tissue barrier integrity. Although commercial TEER meters are available, they are expensive and difficult to customize, which hinders researchers hoping to incorporate them in other research platforms. In the past few years, microcontrollers have risen in popularity for electrical signaling and general programming, of which Arduino is the most popular platform due to its scalability, simplicity and low price. This work presents the development of a completely customized, user-friendly and low-cost TEER meter that is Arduino-based and capable of continuous measurements and automated data collection. We demonstrate the stability of the instrument to measure long-term real-time barrier formation and disruption of an epithelial and endothelial cell line. The design simplicity and low-cost of the components make this technology transferrable to other laboratories for TEER and biological barriers research.

## 1. Introduction

Biological barriers, composed primarily of epithelial and endothelial cells, are multicellular structures that precisely regulate the movement of ions, biomolecules, drugs, cells, and other organisms (Adil et al., 2021; Yeste et al., 2018). A primary characteristic of endothelial and epithelial cells is their ability to create polarized and tight barriers due to the presence of numerous tight junctions (TJs), that allow ions, solutes, and cells to flow specifically through the paracellular space, and adherence junctions, which allow cells to interact with each other (Gonzàlez-Mariscal et al., 2005; Adil et al., 2021; Yeste et al., 2018). Barrier integrity is vital for the physiological activities of the human body (Anderson and Van Itallie, 2009; Krishnakumar et al., 2023). Similarly, many diseases are accompanied by disruption and dysfunction of barrier-forming tissues (endothelial barrier in neurodegenerative disorders, such as Alzheimer’s or Parkinson’s disease (Fang et al., 2023); epithelial barrier in intestinal dysfunctions, including inflammatory bowel disease or immune system (Catalioto et al., 2011)). It is therefore crucial to understand barrier function and properties when studying diseases and developing drugs and treatments (Deli et al., 2005; Nazari et al., 2023; Solovev et al., 2024; Zoio et al., 2021). Thus, physiological and pharmaceutical research has extensively examined the permeability of endothelial/epithelial cell monolayers.

In determining barrier properties, immunostaining for tight junction markers (e.g., Zonula Occludens (ZO) and Occludin) is the gold standard. However, this method has several drawbacks, including time-consuming fixation and staining processes, expensive materials/reagents, and, importantly, does not allow continuous monitoring of cells in real-time. As an alternative to invasive and optical methods for measuring barrier function, electrical measurements can be performed on living epithelial or endothelial cells. In fact, epithelial or endothelial cell layers can be modelled as an electrical parallel between a resistance and a capacitor, which respectively represent the paracellular pathway related to tight junctions (TJ) and the transcellular pathway characterized by the hydrophobic phase of the cell membrane’s capacitance (Cacopardo et al., 2019). An electrical resistance across a monolayer of cell is measured as the transendothelial/epithelial electrical resistance (TEER) (Jones and Chen, 2020; Srinivasan et al., 2015). The measurement principle is based on the application of a direct current (IDC) and on the recording of the resultant voltage (V), giving information on the ohmic resistance of the cellular layer (R = V/IDC). This TEER measurement correlates the electrical properties of an epithelial or endothelial layer with biological aspects such as cell layer confluency and thickness, TJs formation and morphology (Nicolas et al., 2021). Loosely packed cells have lower resistance values because the charge blockage is reduced, whereas tightly packed cells have higher resistance values due to the barrier that prevents charge exchange (Jones & Chen, 2020; Odijk et al., 2015; Yuan et al., 2020). In fact, TEER redout is extensively used in *in vitro* models, as the method does not interfere with cultured cells during operation, provides short response times and good S/N ratios, making it a suitable method for evaluating the effects of drugs, toxins and other substances on the integrity of barriers and studying the mechanisms underlying barrier dysfunction in various diseases (Lucchetti et al., 2024; Rauti et al., 2021; Rauti et al., 2023). Although commercial TEER meters are available, they are expensive (Thousands of €) and difficult to customize, which hinders researchers hoping to incorporate them in other research platforms. The fact that TEER detection is indispensable in numerous research activities makes it essential to develop new technologies that are low-cost, simple, and customizable.

In the past few years, microcontrollers have risen in popularity for electrical signaling and general programming, of which Arduino is the most popular platform due to its scalability, simplicity (open-source programming), and low price (Kondaveeti et al., 2021; Theile et al., 2019). Arduino has indeed found applications, both for research and education, in fabricating and controlling instruments in the scientific community (Chen et al., 2017; Jia et al., 2019). Recently, an Arduino-based instrument capable of measuring TEER in cell cultures has been reported (Jones & Chen, 2020). However, because the electrodes are directly connected to the 5-Volt channel of Arduino, applying a continuous stimulus with the same polarity to the cell culture might lead to the formation of an electrical double layer on the electrodes, thereby damaging the cultured cells.

In this work, we address the challenge of developing a completely customized, user-friendly and low-cost instrument that allows stable TEER measurements across multiple days. More specifically, we describe the construction of a TEER meter that is Arduino-based and capable of continuous measurements and automated data collection (**Figure 1**). Calibration of the electrodes and cell-based experiments were conducted to validate the instrument’s accuracy. As a proof-of-concept, we used TEER electrodes to measure real-time barrier formation and disruption of an epithelial and endothelial cell line. The design simplicity and low-cost of buying the required components make this technology transferrable to other laboratories for TEER research.

**Figure 1.**
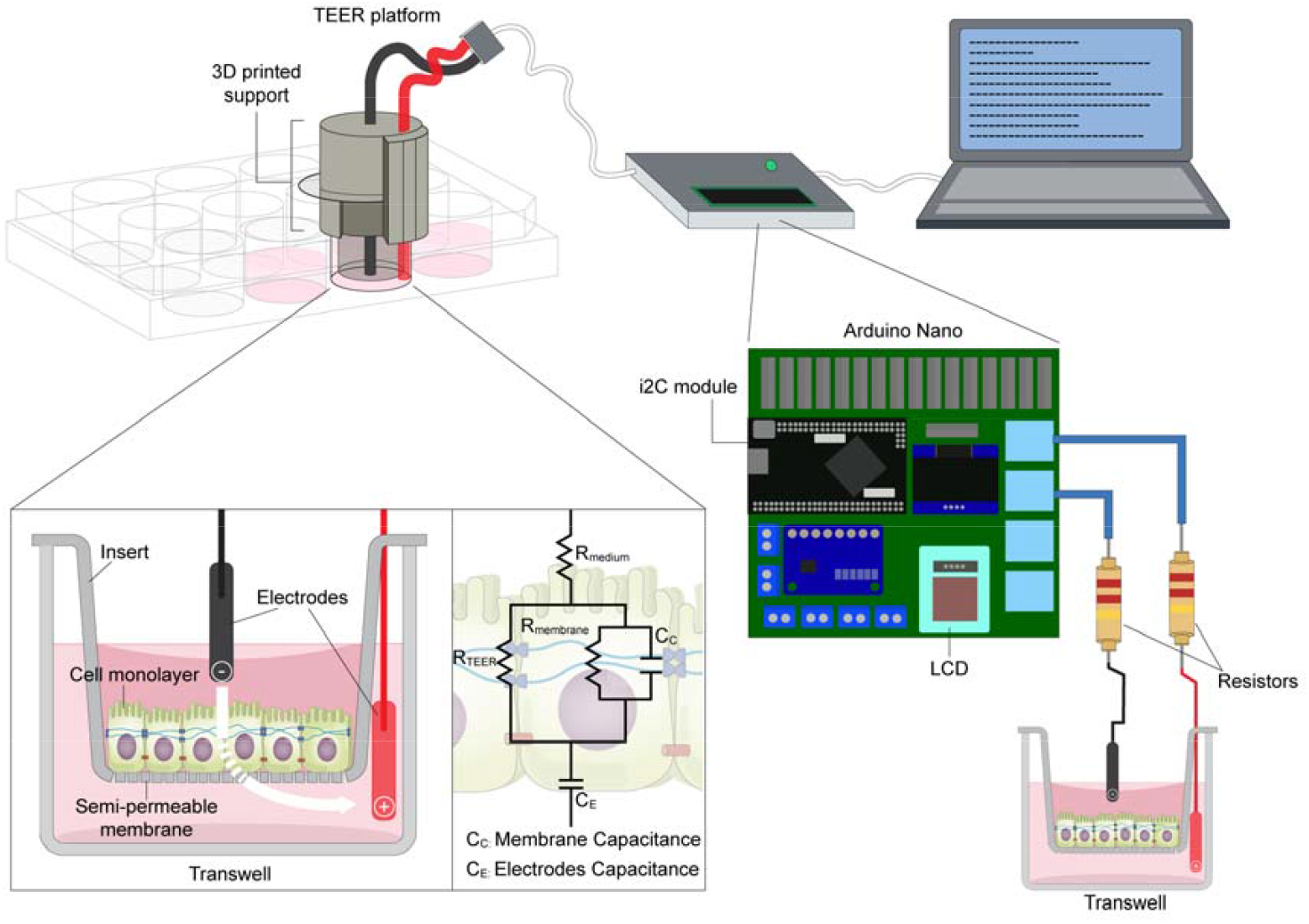
Schematic of TEER Platform. Sketch representing the 3D-printed TEER device connected to the Arduino board.

## 2. Experimental Sections

### 2.1 TEER Device Fabrication

#### 2.1.1 Arduino Board Description

Arduino boards serve as compact programmable devices utilized for constructing electronic projects, whether at a hobbyist or professional level. These boards comprise a microcontroller and multiple input/output pins for connecting sensors, actuators, and other components. These boards are available in various sizes and configurations, with the Arduino Uno and Arduino Nano being the most common. The Arduino Uno boasts 14 digital input/output pins, six analog inputs, a 16 MHz quartz crystal, USB connectivity, and a power jack. On the other hand, the Arduino Nano, a more compact version of the Uno, features 22 digital input/output pins, eight analog inputs, and a smaller form factor. Programming both boards is facilitated through the Arduino Integrated Development Environment (IDE), a software tool enabling code writing, compilation, and uploading to the board. The IDE employs a simplified C++ programming language with built-in libraries and examples to simplify the programming process. Arduino board’s digital input/output pins are designed to handle a maximum current of 40 mA and a maximum voltage of 5 V. Analog input pins enable the board to read analog voltage values from external sensors, with the analog channels converting these input voltages to digital values using a built-in analog-to-digital converter (ADC). The ADC resolution is typically 10 bits, resulting in a range of 0 - 1023 for converted digital values. Analog channels typically use a reference voltage of 5 Volts, resulting in a resolution power of 0.048 Volts (5V/1023). By specifying “analogReference(INTERNAL);” in the code, measurements taken by the analog channels reference an internal potential value of 1.1 volts. Consequently, analogReference(INTERNAL) enhances the ADC’s resolution to 0.012 volts, allowing for more accurate measurement of small amplitude signals.

#### 2.1.2 Arduino Code for TEER Measurement

As mentioned before, Arduino was programmed using the C/C++ language within the Arduino IDE development environment. Due to the compact size of the programmable board and the higher number of available digital channels, in this study, we employed an Arduino Nano open-source electronic prototyping board (Arduino Italy) equipped with an ATmega328 microcontroller. The Arduino board was programmed enabling the board to generate square wave stimuli of 50 ms duration from the digital channels and to measure potential variations between the electrodes positioned across the cell cultures through the analog channels. The entire code is shown with a detailed description in the results section.

#### 2.1.3 Sterilization of the TEER Device

The electrodes were placed in 70% ethanol for 15 minutes and then sterilized under UV lamp for 20 minutes.

### 2.2 Cell Culture

To test and validate our TEER platform, we separately cultured endothelial and epithelial cell layers on Transwells (TWs; Corning) with polyester (PET) membrane (0.4 µm pore size) and monitored the cells over time.

#### Endothelial culture

For the endothelial model, Human Umbilical Vein Endothelial Cells (HUVEC, PromoCell) were used. After thawing, the HUVECs were expanded in low-serum endothelial cell growth medium (Promocell), at 37 °C with 5% CO_2_ in a humidifying incubator, and used at passage p2-p5. Cells were grown at 80-90% confluence before being seeded inside the TWs. Before seeding, the TW membrane was treated with Entactin-Collagen IV-Laminin (ECL) Cell Attachment Matrix (Merck) diluted in Dulbecco’s Modified Eagle Medium (DMEM; 10 µg/cm^2^), for 1h in the incubator. Then, the HUVECs, harvested using trypsin/EDTA solution, were seeded inside the TWs at a density of 20.000/cm^2^ and grown for 19 days.

#### Epithelial culture

For the epithelial model, we used human epithelial colorectal adenocarcinoma cells (Caco-2 cells, ATCC^®^). The passages of the Caco-2 cell line ranged from 26^th^ to 40^th^. After thawing, the Caco-2 cells were cultured routinely in DMEM (Gibco), supplemented with 10% heat-inactivated Fetal Bovine Serum (FBS, Gibco), 1% Glutamax (Gibco), and 1% Penicillin–Streptomycin–Amphotericin B (Anti-anti, Gibco) solution, at 37 °C with 5% CO_2_ in a humidifying incubator. Cells were grown to 80-90% confluency in a T-75 cm^2^ plastic culture flasks (Sarstedt) before being harvested with trypsin/EDTA solution (Merck). Cells were then seeded inside the TWs at a density of 200.000/cm^2^ and grown for 26 days, changing the medium every 3 days of cell culture.

### 2.3 TEER Measurements

The barrier properties of the endothelial and epithelial monolayer were evaluated by our newly developed TEER instrument. TEER values (Ω cm^2^) were calculated and compared to those obtained in a Transwell insert without cells, considered as a blank, in three different individual experiments.

### 2.4 Barrier Disruption

EGTA (ethylene glycol-bis(β-aminoethyl ether)-N,N,N′,N′-tetraacetic acid; Sigma-Aldrich) was prepared at 5 mM with dH_2_O; pH was titrated to 7.4 with HCl 1 M. The medium in the TWs was entirely replaced with the solution containing EGTA and TEER was measured after 10, 20, 30 and 40 minutes.

### 2.5 Immunocytochemistry

Caco-2 and HUVEC cells were rinsed in Phosphate Buffered Saline (PBS) and fixed in 4% paraformaldehyde for 20 min at Room Temperature (RT). Immunocytochemistry was carried out after permeabilization with 0.1% Triton X-100 (Sigma-Aldrich) in PBS for 10 minutes RT and blocking for 30 min with 5 % FBS in PBS. The primary antibodies, anti-β-catenin and VE-cadherin (1:200; Cell Signaling) were applied overnight in PBS and 5% FBS, at 4 °C. Cells were then washed in PBS three times and then incubated with the secondary antibody, anti-rabbit Alexa Fluor-594 (1:500; Invitrogen) in 5% FBS in PBS for 2 hours at RT. After two washes with PBS, cells were incubated with DAPI (1:10.000 in PBS; Cell Signaling) for 10 min at RT to stain the nuclei. After two washes in PBS, imaging was carried out using an inverted epifluorescence microscope (Nikon Eclipse Ti2), with appropriate filter cubes and equipped with 20x/0.50 NA and 40x/0.75 NA objectives. Image reconstruction and processing were done using the open-source imaging software Fiji (Imagej; Schindelin et al., 2012).

### 2.6 Statistical analysis

The results are presented as mean ± SEM, unless otherwise indicated. Statistically significant differences among multiple groups were evaluated by statistics with one-way ANOVA, followed by the Tukey’s test for multiple comparison (GraphPad Prism 8.4.3). The difference between the two data sets was assessed and p < 0.05 was considered statistically significant.

## 3. Results and Discussion

### 3.1 TEER Platform Design and Fabrication

The TEER platform was designed and developed in order to ensure stability during the measurements. It contains two integrated electrodes, constructed using a 0.5 mm diameter silver wire. The electrodes were positioned and secured on a 3D-printed resin support, allowing proper placement within the TW each time (**Figures 2a and 2b**). Using this support was beneficial in reducing variability in TEER readings caused by improper electrode placement. TEER measurements are then continuously recorded using a laptop connected to the platform (**Figure 2c**). We have designed our TEER system to be small and portable (**Figure 2c**) so it can be placed under a biological hood or inside an incubator for cell cultures.

**Figure 2.**
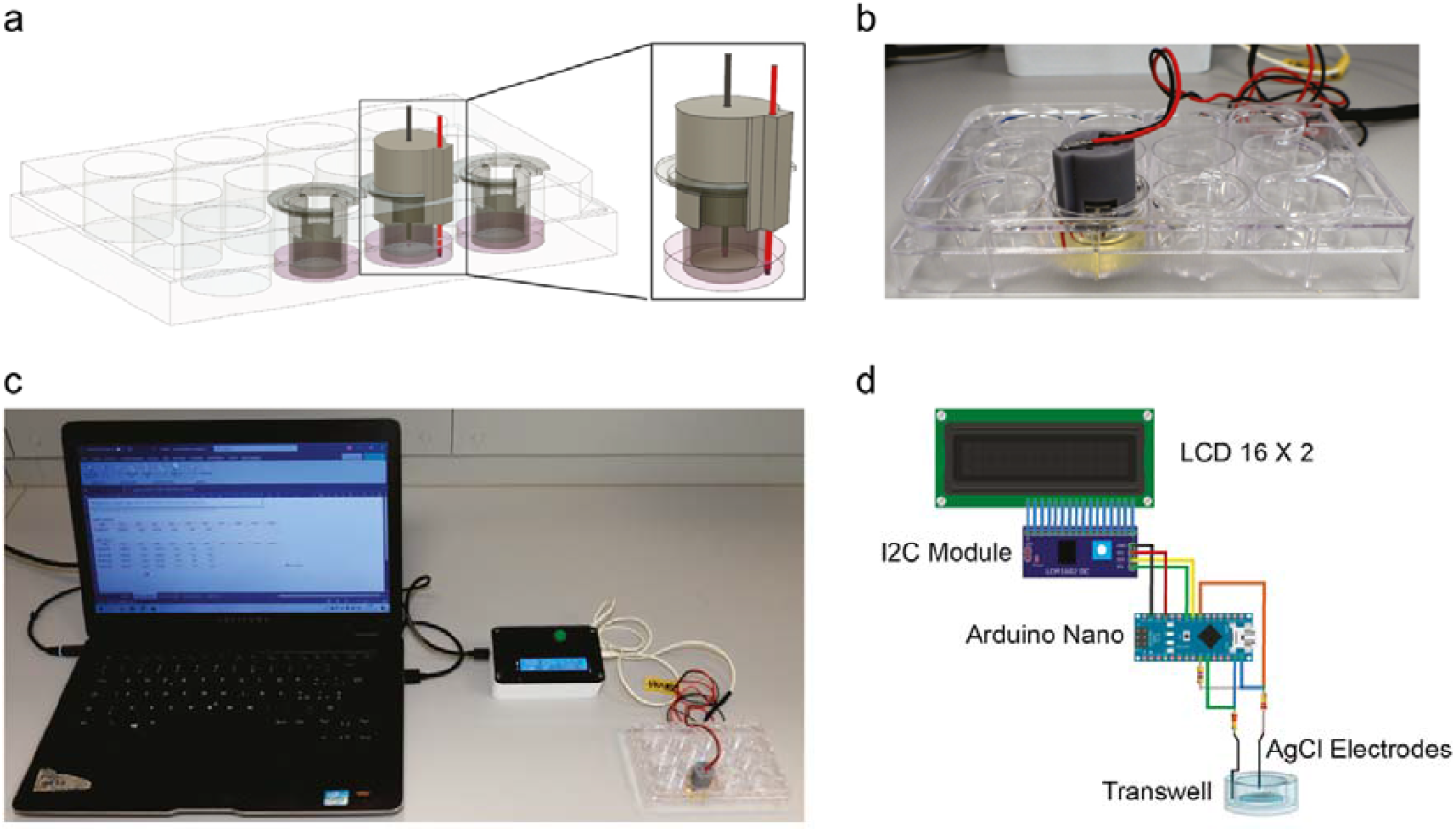
TEER Platform Design. a. A 3D schematic illustration showing the TEER electrodes inserted in a 3D-printed electrode holder and correctly positioned within a Transwell. b. Photograph of the TEER sensor. c. Photograph of the complete TEER measurement setup connected to a PC for data acquisition. d. A diagram illustrating the modules and electronic components comprising the custom-built instrument.

The length of the electrodes was chosen so that the central electrode (located inside the TW) was approximately 2 mm shorter than the lateral electrode (positioned outside the TW). The electrodes were covered for almost their entire length with an insulating sheath. Only the terminal portion of the electrodes (approximately 0.5 mm) was devoid of insulation (**Figure 2a**). This precaution was crucial to prevent varying levels of cell culture medium in different TWs from introducing variability in TEER measurements. The terminal portion of the electrodes was regularly chlorinated (weekly) to ensure stability in measurements over time.

### 3.2 TEER Circuit and Programming

Based on our designed external circuit, each Arduino Nano can read TEER values from 3 pairs of silver chloride electrodes. For simplicity, in this paragraph, we will describe the circuit that allows the Arduino Nano board to read TEER values from a single pair of electrodes. The circuit is detailed in **Figures 2d and 3a**. The external electrode was connected to digital channel D12 through a series resistor R1 (220 kΩ). Between R1 resistor and the silver chloride electrode, digital channel D6 (with a series resistor of 47 Ω) and analog channel A3 were connected. The internal electrode was connected to digital channel D11 through a series resistor R2 (220 kΩ). Between R2 resistor and the electrode, digital channel D7 was connected. Resistors R1 and R2 were used to limit the current passing through the specimen to avoid damaging the cultured cells.

Digital channel D6 was used to discharge the capacitor formed by the silver electrodes and the cellular membranes of the biological specimen after the square wave stimulus was used to measure TEER. In our case, the 47-Ω resistor could also be eliminated since the capacitive currents generated are of low magnitude and would not cause damage to the digital pin D6.

Analog channel A3 was used to calculate the current that flow into the circuit by measuring the potential difference generated by the stimulus applied to the electrodes through digital channel D12. We employed analogReference(INTERNAL) since the currents that pass through the electrodes and, consequently, the biological specimen must remain within a range of a few microamperes, giving rise to potential variations of only a few millivolts. Digital channel D7 was used as a substitute for the ground channel during stimulation originating from digital channel D12 by setting the pin as OUTPUT in the LOW state. After stimulation with digital channel D12, digital channel D11 generated a stimulus of the same duration by setting digital channel D6 as ground (OUTPUT mode, LOW state). As evident from the circuit described in **Figure 3a**, the ground of Arduino is not used. Instead, the ground points are represented alternately by digital channels D7 and D6. This workaround was employed to achieve a polarity reversal of the stimulation electrodes. As described in the experimental results below, the polarity reversal of the electrodes was crucial to prevent electrical double-layer formation on the electrodes, thereby altering their characteristics, such as resistance and capacitance. The second significant advantage of using a bipolar stimulus is preventing electrolytic phenomena harmful to cultured cells.

**Figure 3.**
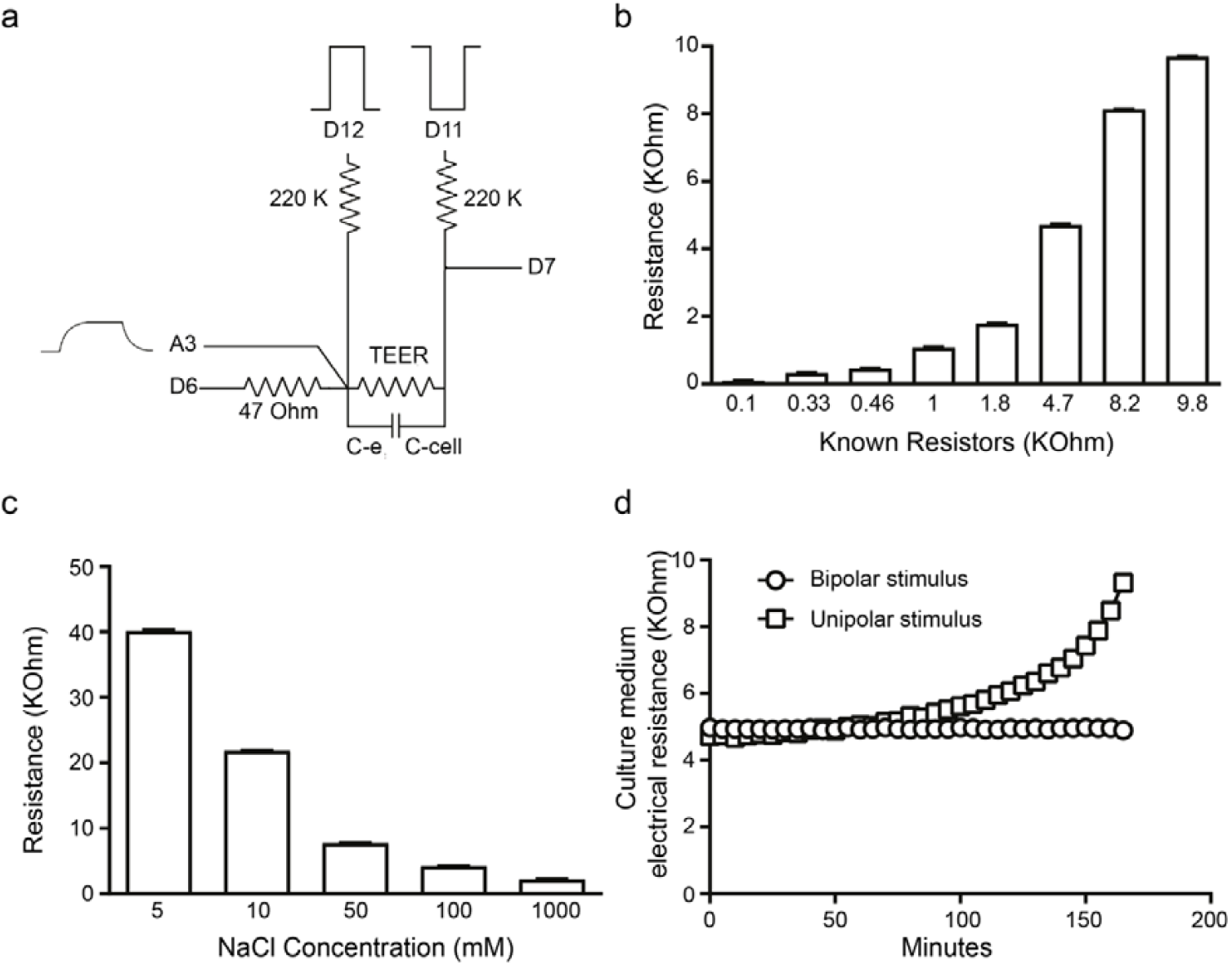
TEER Characterization. a. Schematic of the circuit including digital channels (D11, D12) for generating square wave stimuli, digital channels (D6, D7) for capacitor discharge, and an analog channel (A3) for measuring potential changes induced by the square wave stimuli. The circuit also incorporates 220 kΩ resistors to reduce current levels across the circuit during each stimulus. Additionally, it includes the resistance due to the cells (TEER), the capacitance of the cellular layer (C-cell), and the capacitance of the electrodes (C-e). b-c. Plots representing the measurements of known electrical resistances connected to our custom-built instrument and solutions with varying NaCl concentrations. d. Plot showing the effect of bipolar versus unipolar stimuli on the electrical resistance of electrodes immersed in an electrolyte solution.

As mentioned before, the program was written in Arduino software to measure TEER using two AgCl electrodes and display the results on a Liquid Crystal Display (LCD; **Figure 2c**). In what follows, we will describe the code used in our system.

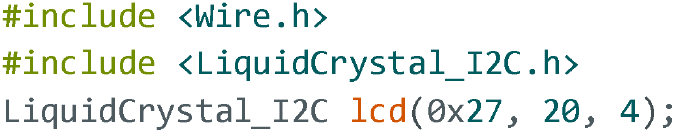

These lines include necessary libraries for I2C communication and for controlling a Liquid Crystal Display using I2C connection. An object lcd of the class LiquidCrystal_I2C is created. This object is used to control the LCD. The parameters passed to the constructor are the I2C address (0x27), number of columns (20), and number of rows (4) of the LCD.

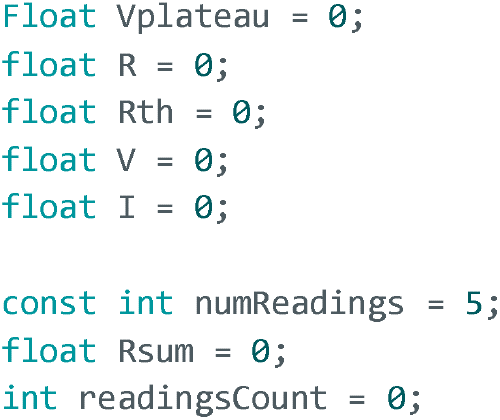

These lines declare several floating-point variables (Vplateau, R, V, i) that will be used for subsequent calculations. Furthermore, this section defines constants and variables related to resistance measurements. numReadings is set to 5, indicating the number of readings to be averaged for a more stable result. Rsum accumulates the sum of resistance readings, and readingsCount keeps track of the number of readings taken.

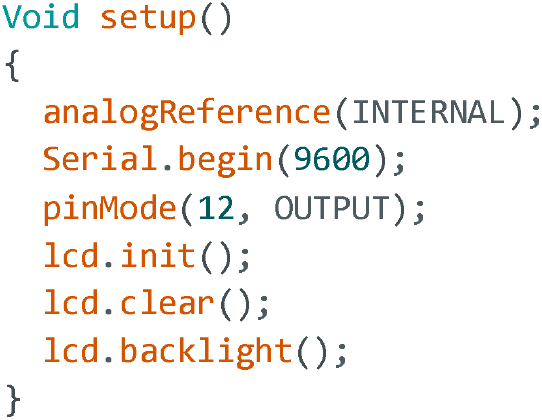

This setup() function is responsible for initializing various settings and preparing the system for operation. The analog reference voltage is set to the internal reference voltage of the Arduino. The INTERNAL reference voltage is typically 1.1V on most Arduino boards. The serial communication at a baud rate of 9600 is initialized. It allows communication between the Arduino and an external device (like a computer) through the serial port. This is useful for debugging and sending data to a computer for monitoring. The pin 12 is set as an output.

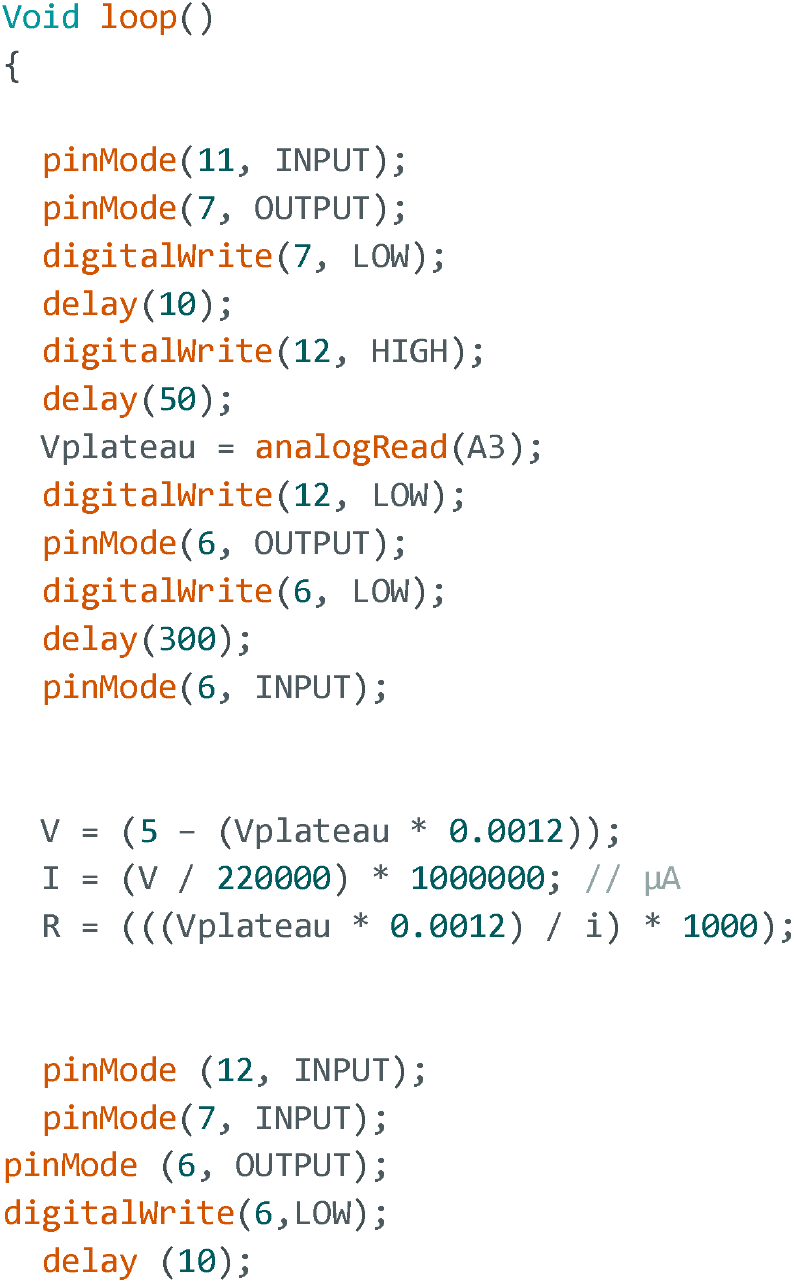
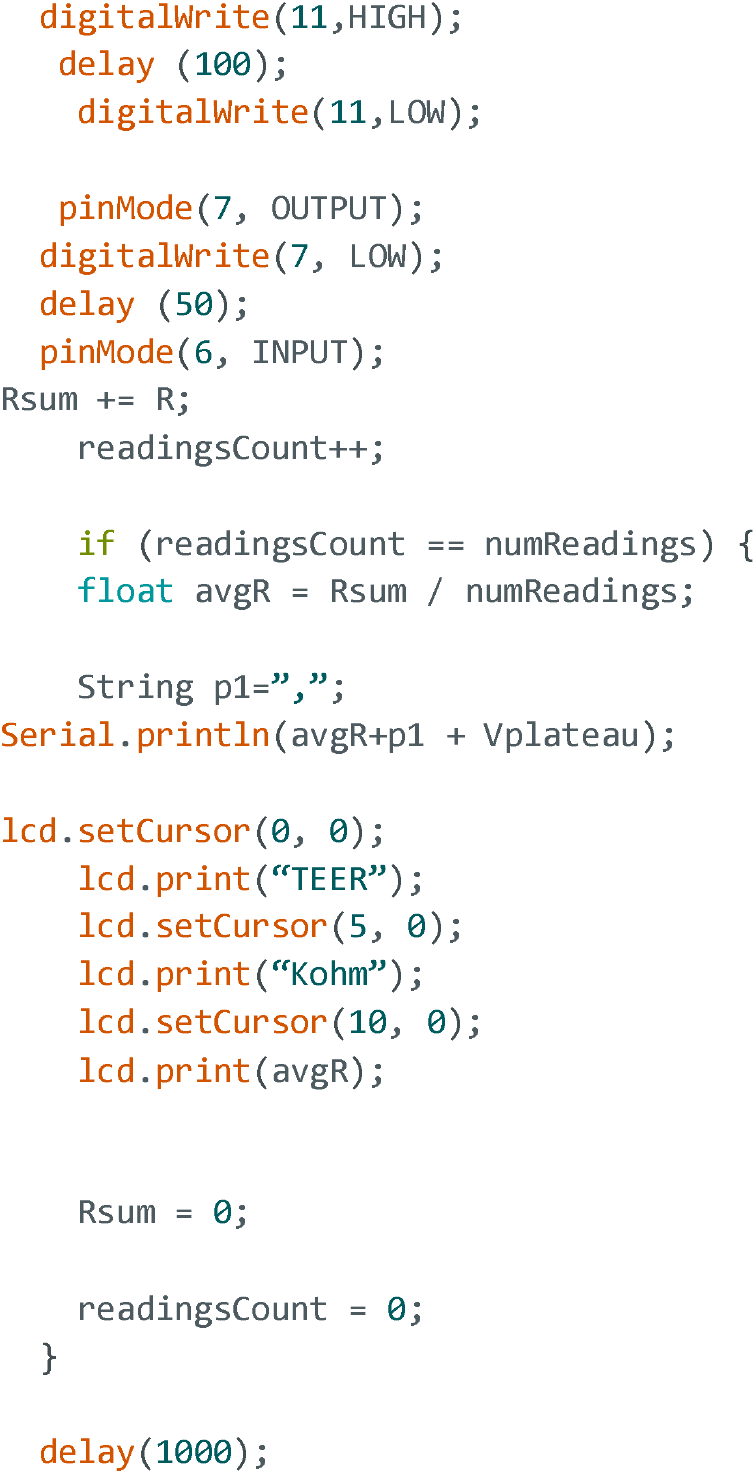

In void loop function, Pin 11 is declared as an input to increase resistance and temporarily exclude it from the circuit. Pin 7 is declared as an output in a low state to use it as a ground momentarily. Pin 12 is set to a high state for 50 ms to allow current to flow between the electrodes for TEER measurement. At the end of the stimulus, the potential difference (Vplateau) between resistor R1 and the electrode is read through analog channel A3, after which pin D12 is set to a low state. Pin 6 is now set as an output in a low state to enable the discharge of capacitors (electrodes and cellular membranes). After a delay of 300 ms, sufficient for complete capacitor-discharge, pin D6 is again set as an input for the next cycle. The following formulas calculate, in order: 1) the potential difference in Volts across resistor R1; 2) the current in microamperes flowing through the circuit after 50 ms from the start of the stimulus when capacitive currents have terminated, leaving only resistive currents; 3) the TEER value by using the calculated values of V and i.

In the following part of the code, the digital channels are configured to invert the stimulation polarity of the electrodes, using pin D11 as the stimulation pin and pin D6 as the ground. The stimulation duration has been set to 100 ms based on experiments conducted by immersing silver chloride electrodes in a culture medium and recording the resistance over several hours. The 100 ms stimulus duration with inverted polarity ensured a stable reading of the medium resistance, avoiding the accumulation of charges on the electrodes (electrical double layer). After stimulation, Pin D7 is set as an output in a low state to enable the discharge of capacitors (electrodes and cellular membranes).

After completing the electrode stimulations, the code accumulates the computed resistance ® in the variable Rsum while incrementing a counter (readingsCount). Upon reaching a specified number of readings (numReadings), it calculates the average resistance (avgR) and outputs the average resistance and the measured voltage to the serial monitor and LCD. The last line of code introduces a delay that determines the frequency of TEER measurements on the sample.

### 3.3 TEER Device Validation and Characterization

Before conducting tests on the self-built instrument, we simulated the operation of the circuit and the installed Arduino sketch using the online Tinkercad Circuits software. During the simulations, we measured a virtual resistance in the range of 0.1 – 10 kΩ connected in parallel with capacitors in the 1-1000 nF range. The simulations allowed us to correctly set the square wave duration output from the Arduino’s digital pins to fully charge the capacitor in parallel before measuring the resistive current flowing through the electrodes. The potential value read by the analog channel A3 exhibits a typical charge and discharge pattern of a parallel RC circuit. The rising phase follows an exponential trend and reaches a plateau when the capacitor is fully charged (**Figure 3a**). At this point, measuring the potential value to calculate the resistive current flowing through the electrodes becomes possible. As hypothesized, increasing the value of the capacitor in parallel required an extension of the square wave duration to achieve a plateau value on channel A3. A stimulus duration of 50 ms was sufficient to obtain a plateau value even with capacitors of 1500 nF.

After the simulations, we tested the circuit constructed on a breadboard with resistors and capacitors identical to those simulated with the software. **Figure 3b** displays the measured resistance values using our homemade instrument with a parallel capacitor of 1000 nF.

To further validate our resistance measurement system, we measured the resistance of distilled water with various concentrations of NaCl (**Figure 3c**), as described in the work of Cacopardo et al., 2019. The resistance was measured by introducing silver electrodes into the saline solution, with a distance of 9 mm between the electrodes. As hypothesized, the measured resistance decreases with an increase in the concentration of NaCl.

As a final step, before utilizing our instrument for measuring TEER on cell cultures, we assessed the culture medium resistance over extended periods (3-4 consecutive hours) to verify the stability of the measured values. As expected, employing unipolar square wave stimuli, the measured resistance values when inserting electrodes into the culture medium did not reach stable levels over time, exhibiting a continuous increase in electrical resistance. This persistent rise in resistance with unipolar stimulation was attributed to charge accumulation on the electrodes, forming the electrical double layer. By employing bipolar stimulation as described previously in the code section, the culture medium resistance values reached stability within a few minutes (electrode conditioning in the culture medium) and remained stable indefinitely (**Figure 3d**).

### 3.4 TEER Measurements to Characterize Barrier Functionality

Once the TEER platform was characterized, we sought to demonstrate its potential and validation by identifying changes in tight junctions of the endothelial and epithelial barrier tissues model. Therefore, we conducted several experiments in which we seeded human endothelial (**Figure 4**) and epithelial (**Figure 5**) cells inside the upper chamber of a TW support. In all experiments, TEER measurements were taken at least 4 times at consistent points, about a couple of minutes after taking the TW out of the incubator. The four measurements were conducted to mitigate the likelihood that the TEER measurements would be influenced by fluctuations in the medium temperature and pH (Srinivasan et al., 2015). Moreover, it demonstrated the stability of the instrument.

**Figure 4.**
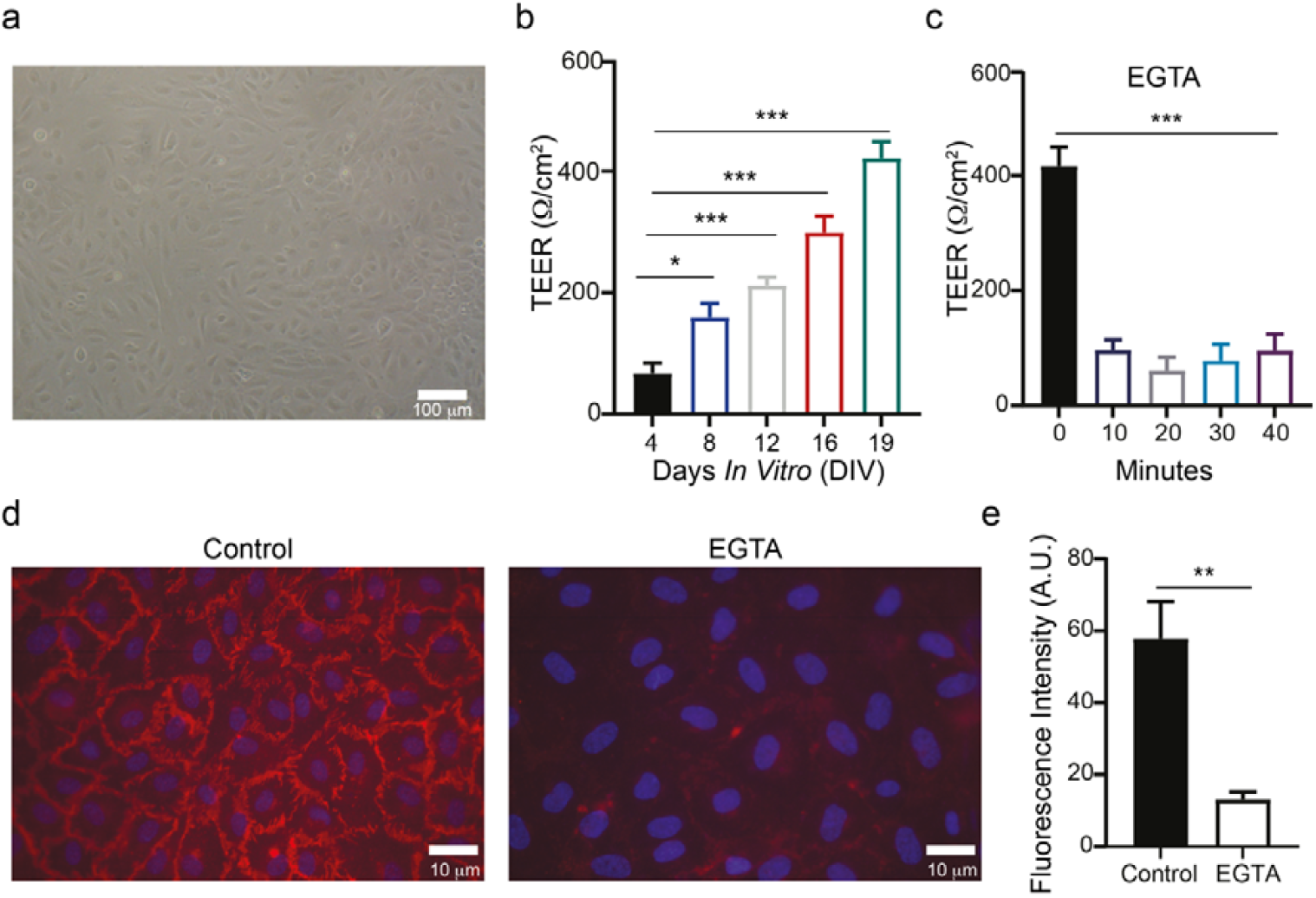
TEER measurements in endothelial cultured cells. a. BF image of cultured HUVEC endothelial cells. b. Plots showing TEER measurements of HUVEC cells in TW insert at different time-points. c. Changes in endothelial barrier function as a result of 5mM EGTA drug application. d. Epifluorescence microscope image reconstruction of VE-Cadherin (red) and DAPI (blue) expression in control and EGTA-treated HUVEC cells. e. Analysis of the VE-Cadherin expression levels from the images presented in panel d.

**Figure 5.**
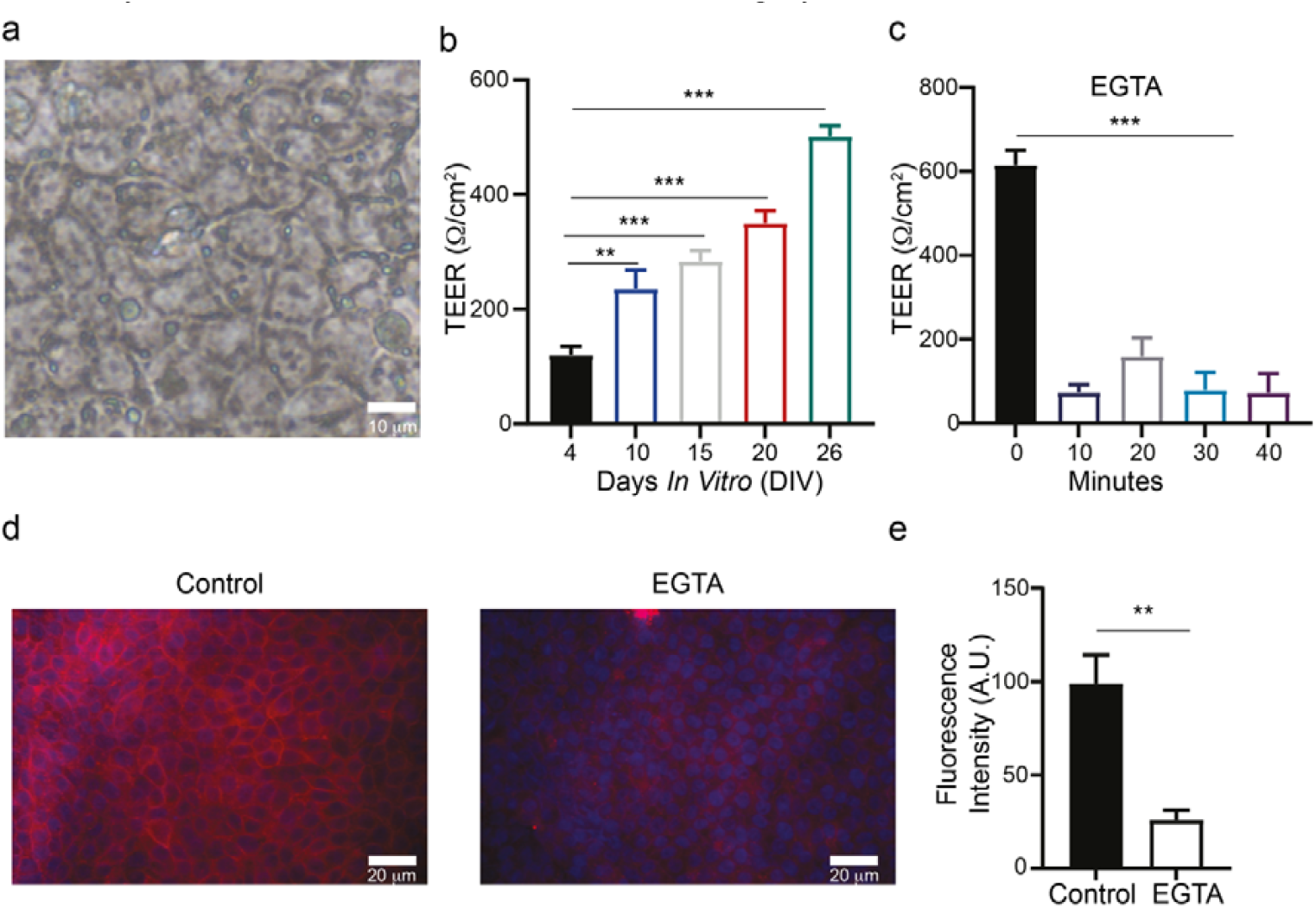
TEER measurements in epithelial cultured model. a. BF image of cultured Caco-2 epithelial cells. b. Plots showing TEER measurements of Caco-2 cells in TW insert at different time-points. c. Changes in epithelial barrier function as a result of 5mM EGTA drug application. d. Epifluorescence microscope image reconstruction of β-catenin (red) and DAPI (blue) expression in control and EGTA-treated Caco-2 cells. e. Analysis of the β-catenin expression levels from the images presented in panel d.

#### 3.4.1 TEER Measurements in an Endothelial Barrier Model

To test our TEER platform, we used Human Umbilical Vein Endothelial Cells (HUVECs, **Figure 4a**) that represent a valid model of endothelial cellular barrier (Motawe et al., 2020; Sultani et al., 2016; Rauti et al., 2021; Rauti et al., 2023). To validate our TEER, endothelial cells were cultured for different days in 12-well plate TWs PET membrane (0.4 µm pore size) until they reach a confluent layer and barrier properties were monitored over time. Due to the small size, the entire setup (electrodes, device and laptop, **Figure 2c**) was inserted inside the biological hood. Once the 3D-printed stage containing the electrodes was placed in the TW, the serial reading was started in the Arduino software and the data were saved on an excel file. The averaged TEER values for the different days *in vitro* are shown in **Figure 4b**. As expected, when comparing HUVEC cells, significant differences were found in TEER values as long as cells were forming a confluent layer, indicating the integrity of the cellular barrier. As shown in Figure **4b**, the TEER of the endothelial monolayer significantly increased (P [ 0.05, one-way ANOVA) from 67.39 ± 16.39 Ω/cm^2^ after 4 days in culture, to 158.8 ± 22.68 Ω/cm^2^ after 8 days in culture, to 210.5 ± 13.91 Ω/cm^2^ after 12 days in culture, to 297.6 ± 26.87 Ω/cm^2^ after 16 days in culture, to 419.5 ± 27.55 Ω/cm^2^ after 19 days in culture (2 samples for each condition, n=3 independent series of culture). To demonstrate the capacity of our TEER to indicate barrier functionality, we monitored changes in permeability due to exposure to EGTA, which is known to damage barrier integrity. When the cell layer reached confluency (as determined by imaging and TEER measurement), 5 mM EGTA in cell culture medium was added to the TW to stress the cell and cause damage to the barrier. TEER was measured at different time points, 10, 20, 30 and 40 minutes after EGTA introduction.

As expected, after introduction of EGTA, we observed a significant decrease in the TEER measurements (**Figure 4c**). Indeed, TEER values decreased from 416.7 ± 32.75 Ω/cm^2^ to 96.25 ± 18.15 Ω/cm^2^ after 10 minutes, 61.25 ± 23.29 Ω/cm^2^ after 20 minutes, 78.54 ± 28.38 Ω/cm^2^ after 30 minutes, to 95.58 ± 28.67 Ω/cm^2^ after 40 minutes, confirming the stability of our TEER device.

As next step, we quantified the correspondence between TEER values and the barrier integrity via immunocytochemistry, measuring VE-Cadherin expression, before and after EGTA. To do so, HUVEC cells were immunostained for one of the main tight-junction (TJ) protein stained for VE-Cadherin protein (in red, **Figure 4d**) and DAPI (in blue, **Figure 4d**) for the nuclei. As shown in **Figure 4e**, cells treated with EGTA showed a significant (P [ 0.01, one-way ANOVA) reduction in VE-Cadherin expression (from 57.93 ± 10.23 A.U. before EGTA, to 13.11 ± 2.122 A.U., after EGTA treatment), confirming the ability of our TEER device to measure barrier integrity.

#### 3.4.2 TEER Measurements in an Epithelial Barrier Model

To further validate our TEER platform, we decided to test it on an epithelial cellular barrier model. Therefore, we conducted several experiments in which we seeded human epithelial colorectal adenocarcinoma cells (Caco-2 cells; **Figure 5a**) inside the TW. As for the HUVEC, Caco-2 cells were cultured for different days in 12-well plate TWs PET membrane (0.4 µm pore size) until they reach a confluent layer and barrier properties were monitored over time.

The averaged TEER values for the different days *in vitro* are shown in **Figure 5b**. As expected, when comparing Caco-2 cells, significant differences were found in TEER values as long as cells were forming a confluent layer, indicating the integrity of the cellular barrier. As shown in Figure **5b**, the TEER of the epithelial monolayer significantly increased (P [ 0.001, one-way ANOVA) from 120.2 ± 15.23 Ω/cm^2^ after 4 days in culture, to 235.8 ± 32.33 Ω/cm^2^ after 10 days in culture, to 283.5 ± 18.48 Ω/cm^2^ after 15 days, to 345.7 ± 21.69 Ω/cm^2^ after 20 days, to 501 ± 18.87 Ω/cm^2^ after 26 days in culture (2 samples for each condition, n=3 independent series of culture). Next, we monitored changes in permeability due to exposure to 5 mM EGTA and measured TEER 10, 20, 30 and 40 minutes after EGTA exposure.

As expected, we observed a significant decrease in the TEER measurements (**Figure 5c**). Indeed, TEER values decreased from 615 ± 35.06 Ω/cm^2^ to 73.33 ± 19.24 Ω/cm^2^ after 10 minutes, 153.2 ± 39.32 Ω/cm^2^ after 20 minutes, 79.17 ± 42.74 Ω/cm^2^ after 30 minutes, to 72.92 ± 45.09 Ω/cm^2^ after 40 minutes, confirming the stability of our TEER device.

As next step, we quantified the correspondence between TEER values and the barrier integrity via immunocytochemistry, measuring β-Catenin expression, before and after EGTA. To do so, Caco-2 cells were immunostained for β-Catenin protein (in red, **Figure 5d**) and DAPI (in blue, **Figure 5d**) for the nuclei. As shown in **Figure 5e**, cells treated with EGTA showed a significant (P [ 0.01, one-way ANOVA) reduction in β-Catenin expression (from 99.08 ± 15.25 A.U. before EGTA, to 26.10 ± 5.16 A.U., after EGTA treatment), confirming the ability of our TEER device to measure barrier integrity.

## 4. Conclusions

In this work, we developed a completely customizable and low-cost 3D-printed TEER device to measure endothelial and epithelial cellular barrier properties. Specifically, we report the design, fabrication and validation of a TEER device using Arduino code. A key strength of this work is its ability to enable researchers, even without extensive knowledge of electronics, to reproduce a low-cost instrument that allows precise and stable TEER measurements in different cell cultures. Our system relies on two electrodes that are mounted on a 3D-printed holder that can be completely customized according to the platform used for cell culture. We validated our system in experiments using endothelial and epithelial cells, showing that the TEER measurements are in direct correlation with cell concentration and TJs expression. We further demonstrated the accuracy of our system to identify changes in barrier integrity following exposure to EGTA. It is worth noting that there has been a recent study in which Arduino was used to develop a TEER system (Jones and Chen, 2020). While their approach is similar to ours, in this study, the circuit was designed to achieve electrode polarity inversion to prevent the formation of the electrical double layer and the consequent alteration of the resistance and capacitance values of the electrodes. The second significant advantage of using a bipolar stimulus is preventing electrolytic phenomena harmful to cultured cells.

Additionally, the system’s modularity (fabricated with 3D printing) and small size make it compatible with long-term measurements in cell culture incubators as well as with different cell culture platforms. As a results of its simplicity and customizability, this instrument will greatly benefit scientific research on biological barriers.

### Credit authorship contribution statement

DL designed and conceived the TEER device and wrote the Arduino code. LM performed the cell biology and immunofluorescence experiments and analysis. AS contributed to the TEER measurements and data analysis. DL and RR conceived the study and the experimental design. DL, LM, AS and RR wrote the manuscript.

## Acknowledgements

This work was supported by the European Union - NextGenerationEU within the framework of PNRR Mission 4 - Component 2 - Investment 1.1 under the Italian Ministry of University and Research (MUR) programme “PRIN 2022 PNRR” - grant number P2022TKL5T_001 Calinero - CUP: H53D23009070001 and by the European Union - NextGenerationEU, Mission 4, Component 1, under the Italian Ministry of University and Research (MUR) National Innovation Ecosystem grant ECS00000041 - VITALITY - CUP H33C22000430006.

We are also especially grateful to Caterina Di Pietro for the art work.

## Data availability

The datasets generated for this study are available on request to the corresponding author.

